# Plagues of Desert Locust: No invasion risk to China

**DOI:** 10.1101/2020.03.03.973834

**Authors:** Yun-Ping Wang, Ming-Fei Wu, Pei-Jiong Lin, Yao Wang, Ai-Dong Chen, Yu-Ying Jiang, Bao-Ping Zhai, Jason W. Chapman, Gao Hu

## Abstract

Recently, the most serious upsurge of desert locust (*Schistocerca gregaria*) in the last 25 years is spreading across eastern Africa and southwestern Asia. Parts of the desert locust ‘invasion area’, namely the northern border areas of Pakistan and India are very close to China, and whether locust swarms will invade China is of wide concern. To answer this question, we identified areas of potentially suitable habitat for the desert locust within China based on historical precipitation and temperature data, and found that parts of Xinjiang and Inner Mongolia provinces could provide ephemeral habitat in summer, but these places are remote from any other desert locust breeding area. Presently, the desert locust populations in Pakistan and India are mature and have laid eggs, and are less likely to spread long distances. The next generation of adults will appear in April and May, and so we examined twenty years’ historical wind data (2000–2019) for this period. Our results showed that winds at the height of locust swarm flight blew eastward during April and May, but the wind speeds were quite slow and would not facilitate desert locust eastward migration over large distances. Furthermore, simulated trajectories of desert locust swarms with 10 days’ migration mostly ended within India. The most easterly point of these trajectories just reached eastern India, close to the border between India and Myanmar, and this is very close to the eastern border of the invasion area of desert locust described in previous studies. In conclusion, the risk that the desert locust will invade China is very low.

---

Desert locust, *Schistocerca gregaria*, is one of the most devastating migratory pests in the world [1-4]. It is highly mobile and feeds on large quantities of any kind of green vegetation, including crops, pasture, and fodder [1-4], and even a moderate swarm measuring 10 km^2^ would eat some 1000 tons of vegetation daily. Recently, the most serious upsurge of desert locust since the last serious outbreak in 1994–1995 is spreading across eastern Africa and southwestern Asia. From the figures published by FAO in early February 2020, more than 280,000 ha in 13 countries were infected, and the infected area is still increasing as the locust swarms spread [5]. Among the “Invasion areas’ defined from earlier studies [1], the northern borders of Pakistan and India are very close to China [5]. China is struggling to control fall armyworm (*Spodoptera frugiperda*), a migratory pest that invaded China via India and Myanmar last year [6], and consequently, whether desert locust swarms will invade China is of wide concern.

To answer this question, we first checked whether there are potentially suitable habitats for desert locust within China. Previous studies showed that (i) the desert locust is well-adapted to live in arid and semi-arid habitats where annual precipitation is roughly between 0–400 mm [7], and at least 20 mm of rain falling in a short period (or its equivalent in run-off) is required for egg development [1]; (ii) the air temperature range for egg and hopper (nymph) development is between 20–35 °C [4]; and (iii) if there is more than seven days at 10 °C, egg mortality is considerably increased [8]. Thus, we derived the 20 years’ (2000–2019) mean annual precipitation from the Climate Prediction Center Merged Analysis of Precipitation data, and 20 years’ (2000–2019) mean monthly air temperature from National Centers for Environmental Prediction (NCEP)/National Center for Atmospheric Research (NCAR) reanalysis data. The mean annual precipitation data showed that most areas in South Asia, Southeast Asia and East Asia are too wet for desert locust persistence (annual rainfall ≥ 400 mm), such as most of India, Myanmar and most of China (Fig. 1). In China, only in parts of Xinjiang, Inner Mongolia, Tibet and Qinghai provinces, the annual rainfall is ≤ 400 mm. But the mean air temperature in the Qinghai-Tibet Plateau is still quite cold in July (≤ 20 °C). Further, in most areas of China, the mean air temperature in January is ≤ 10 °C (fig. 1), and this shows that most areas of China are too cold for desert locusts to survive in winter. Taking this information together, only parts of Xinjiang and Inner Mongolia provinces could conceivably be suitable habitat for the desert locust in China, and these would just be ephemeral habitats in summer.

**Fig. 1:**
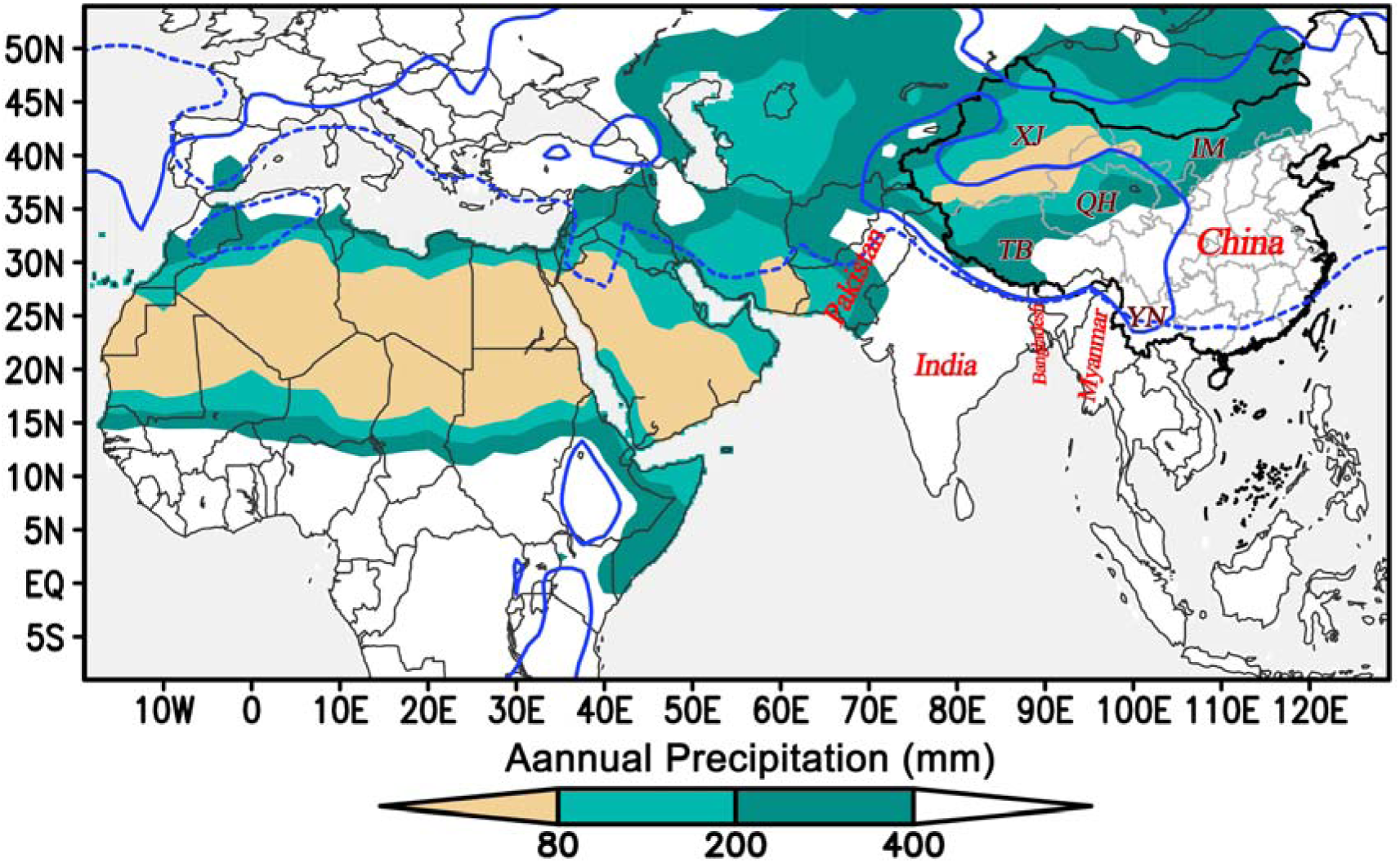
Mean annual precipitation (filled color), 10 °C isotherm in January (blue dotted line) and 20 °C isotherm in July (blue solid line) averaged for 20 years (2000–2019). Potentially suitable habitat for the desert locust should, at least, satisfy conditions with an annual precipitation ≤ 400 mm and air temperature in July ≥ 20°C. As eggs cannot survive when the temperature is below 10°C, for year-round breeding the temperatures in winter should be above 10°C. Taking all these things together, only small areas in Xinjiang and Inner Mongolia provinces could provide an ephemeral (summer) habitat for the desert locust in China. TB-Tibet, YN-Yunnan, QH-Qinghai, IM-Inner Mongolia.

Although we found there are few suitable habitats for desert locust in China (see above paragraph), there might be still a notional threat if the swarms from the border between Pakistan and India enter China. However, the Himalayan mountains exceed 7000 m, and form a natural barrier stopping all or almost all insect migration -- this is particularly likely with desert locusts as the lowest air temperatures (in the absence of sun) at which sustained swarm flight begins is 23-24°C, and even in continuous bright sunshine flight does not begin at air temperatures much below 17°C [1]. Thus, the only possible route that desert locust swarms could move eastward and enter the Yunnan province of China is by crossing India and Myanmar. Gregarious swarms can spread at low-level near the ground or at high-level at hundreds of meters above ground (during strong convection) in daytime, while spend the night roosting in vegetation [2,4]. The low flights often occur during rather cool, overcast weather or in the late afternoon. Locust low fliers can stabilize their ground speed at about 4.0 m/s by varying their air speed and heading direction according to the wind speed, and thus they can move up to 150 km in one day by 10-hr sustained flying [2,4]. The straight distance from the Pakistan-India border to Yunnan province is about 3000 km. Swarms from this region would need 20 days’ sustained flying to cover this distance if they just keep flying at lower level near ground, and this is quite impossible.

The desert locust swarms also can travel over hundreds and thousands of kilometers by riding high-level winds, for example from North-West Africa to the British Isles in 1954 and from West Africa to the Caribbean, a distance of 5,000 km in about ten days, in 1988 [9]. High-flying swarming locusts do not have any single preferred orientation and turn back towards the center of the swarm when they reach the edges in order to maintain swarm cohesion. Thus, the speed and direction of swarm displacement is the same as the wind at the height of flight [1,2,9,10]. So, we then checked whether there are suitable winds at high level (about 1500 m above ground) to carry locust eastwards. Long-range swarm migration is more likely to happen before sexual maturity [2,10]. At this moment, the swarms in Pakistan and India are mature and have laid eggs, and are thus less likely spread over long distances. In this region, the mean temperature in February, March and April in last 20 years is 21.07±0.16°C, 26.93±0.14°C and 32.30±0.10°C, respectively. Based on the relationship between air temperature and the desert locust development [2,4,8], the next generation of locust adults will appear in April and May, after about 30 days as eggs and 30-40 days as hoppers. We thus focused on the period in April and May, and analyzed 20 years’ (2000–2019) historical climatic data derived from NCEP/NCAR reanalysis data. We found that the winds at 850 hPa level (about 1500 m above sea level) blew consistently toward the east during daytime in these two months (Rayleigh test; April: mean value 88 °, *r* = 0.58, P < 0.0001, *n*=600; May: 80 °, *r* = 0.76. P < 0.0001, *n*=620; the *r*-value is a measure of the clustering of the angular distribution from 0 to 1), but the wind speed is quite slow (April: 3.32 ± 0.07 m/s (mean value ± standard error (S.E.)), n=600; May: 4.23 ± 0.08 m/s, n=620) (Fig. 2). As the desert locust cannot fly when the air temperature at flight altitude is below 20 °C [see above, 1,9], we also identified the area of air temperatures at 850 hPa level ≥ 20 °C in late April and late May (Fig. 2). The area with a suitable temperature for locust swarm flight covered most of India, Myanmar and Bangladesh, but in China only Yunnan province was suitable. Therefore, it is conceivable that the desert locust might reach Yunnan province but cannot fly further north in China by windborne migration in April and May.

**Fig. 2:**
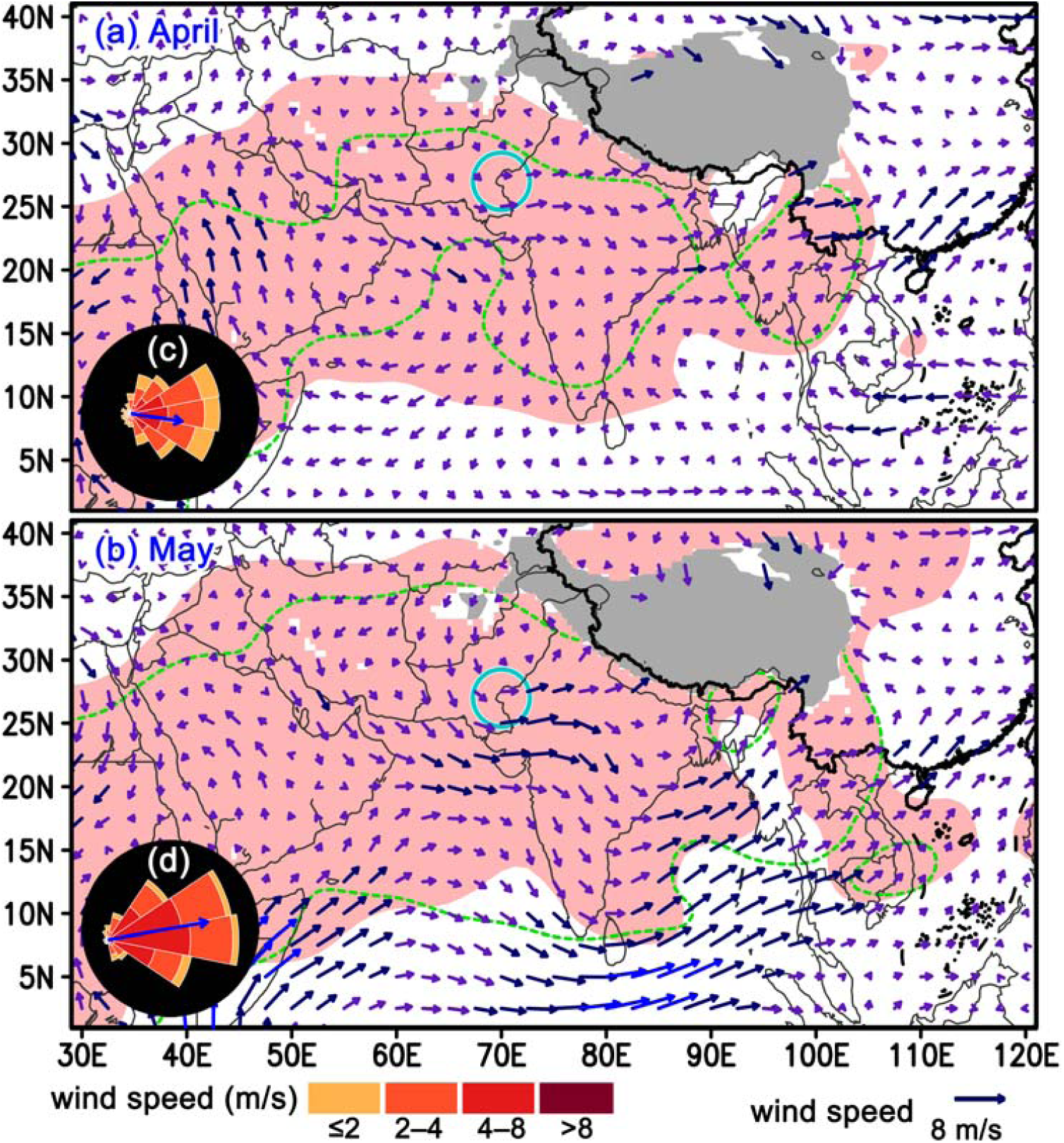
Mean wind (vector arrows) and temperature (the area filled with light red is ≥ 20°C on 30 April or 31 May, and green dashed line is on 20°C isotherm on 1 April or 1 May) conditions at 850 hPa (about 1500 above sea level) in April and May in last 20 years (2000–2019). Subplots in (c) and (d) show the histogram distribution of wind direction and wind speed in the border between Pakistan and India (indicated by blue circles). The grey filled area shows the Qinghai-Tibet Plateau.

Lastly, we modelled the long-distance migration of the desert locust with the Hybrid Single-Particle Lagrangian Integrated Trajectory (HYSPLIT) model of the National Oceanic and Atmospheric Administration (NOAA) [11]. This model is designed for computing three-dimensional trajectories of air parcels and applied extensively to study migratory trajectories of many insect species [12]. Here, we made the following assumptions for the trajectory simulation of the desert locust: (i) the speed and direction of its displacement is the same as the wind at its flight height; (ii) daytime flying occurs mostly from 0900 a.m. (local time, i.e. 0300 UTC) to 1900 a.m.; (iii) locusts cannot fly at air temperatures below 20°C; and (iv) long-distance flight can be sustained for up to 10 days. Because we do not have the meteorological data for April and May in this year (2020), we used the meteorological conditions at flight altitude from the past 5 years (2015–2019). In total, 305 forward trajectories starting from a point at the border between Pakistan and India (27°N, 70°E) were calculated (Fig. 3). Most trajectories went eastwards, and the endpoints were located east of the startpoint (Rayleigh test; April: mean value 90°, *r* = 0.91, P < 0.0001, *n*=150; May: 95°, *r* = 0.94, P < 0.0001, *n*=151). Due to the slowness of the winds, migration distances were quite short (April: 917.3 ± 16.9 km; May: 1116.1 ± 11.5 km), and most trajectories ended within India (Fig. 3). All these results indicate that it is impossible for desert locust to reach China by 10 days’ windborne migration.

**Fig. 3:**
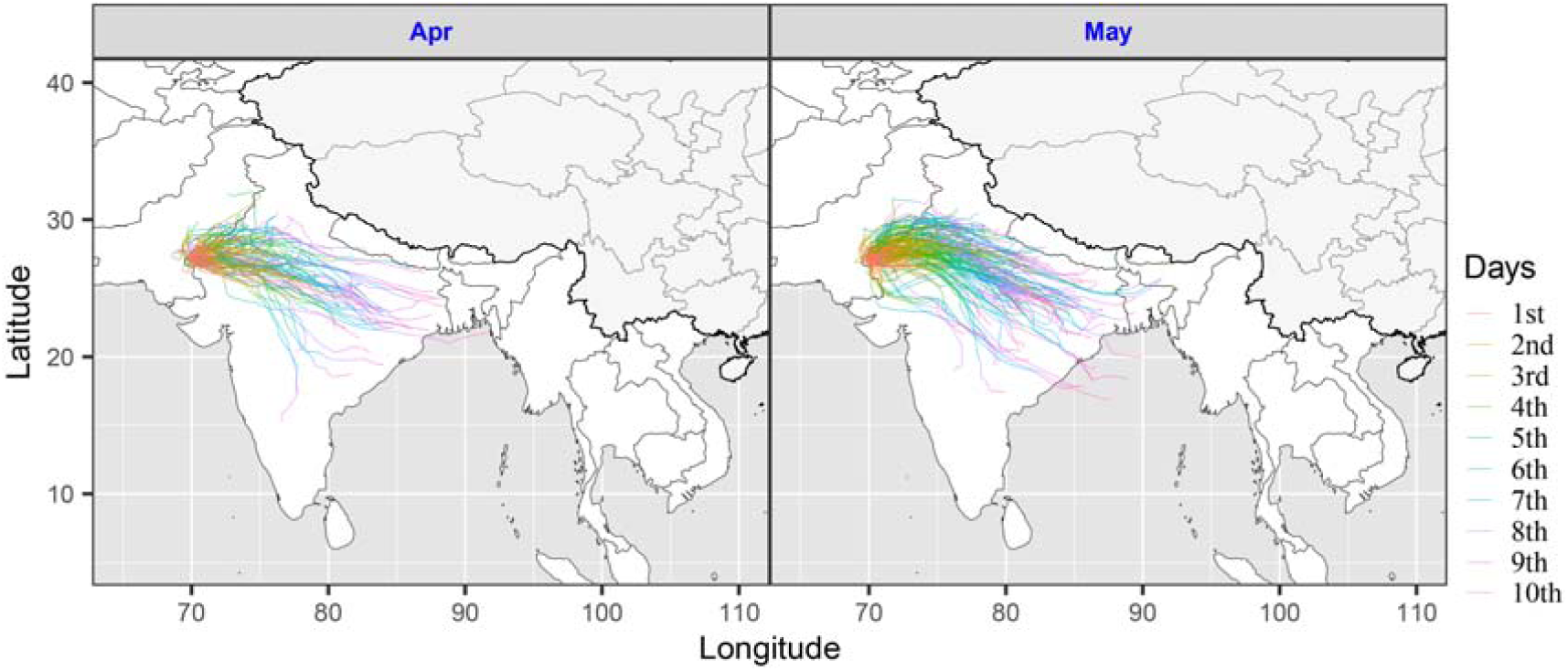
Simulated forward trajectories started from a point at the border between Pakistan and India (27°N, 70°E) in last 5 years (2015–2019). These trajectories were calculated by the Hybrid Single-Particle Lagrangian Integrated Trajectory (HYSPLIT) model.

## Conclusions

The distribution of desert locust is very well known and, during plagues, it can cover the arid and semi-arid region from the Atlantic coast of West Africa to eastern India. The most easterly distribution during invasion periods reaches eastern India and Bangladesh but does not include any part of China [1,4]. The potentially suitable area identified in this study was based on an annual precipitation **≤** 400 mm and mean air temperature in January **≥** 10°, and this area is consistent with the desert locust ‘recession area’, that is, the area occupied when there are few, if any, swarms present [1,4,7]. In China, only parts of Xinjiang and Inner Mongolia provinces could provide ephemeral habitat in summer, and these regions are far away from any other desert locust breeding area. In the past, only one individual of the solitaria form of desert locust was detected in Zhangmu District, Nyalam County in Tibet (location at about 28.33° N, 86°E, elevation; 2250 m) on 29 April 1974, and this place is on the southern slope of the Himalayan Mountains [13]. During April and May, the winds at height of about 1500 m above sea level blow eastward, but the wind speed is quite slow. In our trajectory simulations, most trajectories with 10 days’ migration ended within India, and the furthest just reached eastern India, or close to the border between India and Myanmar. The most easterly point of our trajectories is almost the same as the eastern border of the invasion area of desert locust described in previous studies [4,7]. In conclusion, it is very unlikely that agriculturally-significant populations of the desert locust will invade China.

## Acknowledgements

This work was supported though grants to G.H. by the National Natural Science Foundation of China (31822043), the Natural Science Foundation of Jiangsu Province (BK20170026) and the Fundamental Research Funds for the Central Universities (KJYQ201902).

